# A missense variant effect prediction and annotation resource for SARS-CoV-2

**DOI:** 10.1101/2021.02.24.432721

**Authors:** Alistair Dunham, Gwendolyn M Jang, Monita Muralidharan, Danielle Swaney, Pedro Beltrao

## Abstract

The COVID19 pandemic is a global crisis severely impacting many people across the world. An important part of the response is monitoring viral variants and determining the impact they have on viral properties, such as infectivity, disease severity and interactions with drugs and vaccines. In this work we generate and make available computational variant effect predictions for all possible single amino-acid substitutions to SARS-CoV-2 in order to complement and facilitate experiments and expert analysis. The resulting dataset contains predictions from evolutionary conservation and protein and complex structural models, combined with viral phosphosites, experimental results and variant frequencies. We demonstrate predictions’ effectiveness by comparing them with expectations from variant frequency and prior experiments. We then identify higher frequency variants with significant predicted effects as well as finding variants measured to impact antibody binding that are least likely to impact other viral functions. A web portal is available at sars.mutfunc.com, where the dataset can be searched and downloaded.

## Introduction

A novel coronavirus, SARS-CoV-2, emerged in late 2019 and spread across the world to cause the COVID19 pandemic, which remains a global health emergency. The pandemic has led to many deaths, an even greater number of other medical consequences and a deep and far reaching economic impact. This has made responding to it one of the largest challenges in modern times. This response takes many forms, including drug and vaccine research, public health policy and economic stimulus. A key aspect to the global response is monitoring viral mutations, which allows us to track the virus geographically and follow the emergence of mutations that might lead to impactful changes in the virus. This makes the ability to quickly analyse the potential consequences of new variants invaluable.

Many viral mutations have occurred, including a small number that have reached high frequencies regionally or worldwide, and recently three major viral substrains, B.1.1.7, B.1.351 and P.1, have caused significant concern because of their rapid spread and potential for antibody evasion (Mahase, 2021). The most common form of mutation is substitutions, where one RNA nucleotide changes to another. This impacts both viral RNA structure, known to influence function (Ziv et al., 2020), and protein sequence, which we focus on here. Amino acid substitutions alter protein’s chemistry and can change structure, stability and activity, in turn impacting biological function. While most substitutions have little or no effect others have significant impact. For example the N439K variant in the Spike protein has been found to reduce binding by some antibodies and increase affinity with human ACE2, which is bound during entry to host cells (Thomson et al., 2021). Similarly the E484K Spike variant observed in B.1.351 and P.1 has caused concern because it affects antibody binding (Greaney et al., 2021; Wise, 2021).

Many computational tools have been developed to predict the consequences of mutations based on protein sequence and structure. Sequence is informative because the prevalence of a variant at similar positions across species relates to the likelihood of it being tolerated. This is utilised by tools like SIFT4G (Vaser et al., 2015) and EVCouplings (Hopf et al., 2019). EVCouplings predictions for SARS-CoV-2 are publicly available online (https://marks.hms.harvard.edu/sars-cov-2/). Protein structure is used by tools like FoldX (Schymkowitz et al., 2005) and Rosetta (Kellogg et al., 2011) to model the effects of mutations on thermodynamic properties, including on complex binding if complex models are available. Machine learning is also used to combine feature types (Gnad et al., 2013; Gray et al., 2018). However, running predictors can be technically challenging and computationally expensive, which can be prohibitive for many scientists, preventing them taking advantage of predictions.

The Mutfunc web service (Wagih et al., 2018) was developed in our lab to provide an interface for pre-computed predictions for all possible *H. sapiens*, *M, musculus* and *S. cerevisiae* variants. We apply a similar analysis to the proteins of SARS-CoV-2, combining predictions based on sequence conservation, protein structures, known protein-protein interfaces, phosphosites and observed frequencies. A web interface is available at sars.mutfunc.com, which allows users to search and download the dataset and provides visualisations of structures and alignments. We analyse the resource, validating the use of predictors and showing its benefits for identifying potential variants of interest or concern among higher frequency variants, those in variant strains and those with antibody evasion potential.

## Results

### Mutfunc: SARS-CoV-2 Dataset

We collected sequences and structures of the 28 SARS-CoV-2 proteins from a range of sources (*Figure 1A*). Sequences came from Uniprot (The UniProt Consortium, 2019) and were fed to SIFT4G to make conservation based predictions for all possible variants. SWISS-MODELs SARS-CoV-2 repository (Bienert et al., 2017; Waterhouse et al., 2018) was used to source the highest quality experimental or homology structural models for each protein, using multiple models to cover more positions where possible (see methods). Structural coverage was high for the 19 proteins with structures (*Figure 1B*); only nsp4 has coverage below 50%. These models were used to generate FoldX ΔΔG predictions for all possible variants at positions covered by models. Complex models from SWISS-MODEL and PDBe were analysed with FoldX to identify interface residues and predict the effect on binding energy of all possible interface mutations. Sixteen complex structures were identified at the time of publication (*Figure 1C*), 10 between viral proteins, 2 with human proteins and 4 with antibodies. Viral phosphosites were integrated from Bouhaddou et al. (2020). Variant frequencies across all samples up to February 2021 were calculated from a sequence alignment of over 235,900 public SARS-CoV-2 sequences (Lanfear and Mansfield, 2020; Turakhia et al., 2020). Sequences are dominated by samples from the UK (75.0%) and USA (14.4%), meaning frequencies mostly reflect these regions. Experimental antibody evasion data (Greaney et al., 2021) and some individual variant annotations (Public Health England, 2020; Tegally et al., 2020; Faria et al., 2021; Mahase, 2021; Kemp et al., 2021) complete the Mutfunc: SARS-CoV-2 dataset.

**Figure 1.**
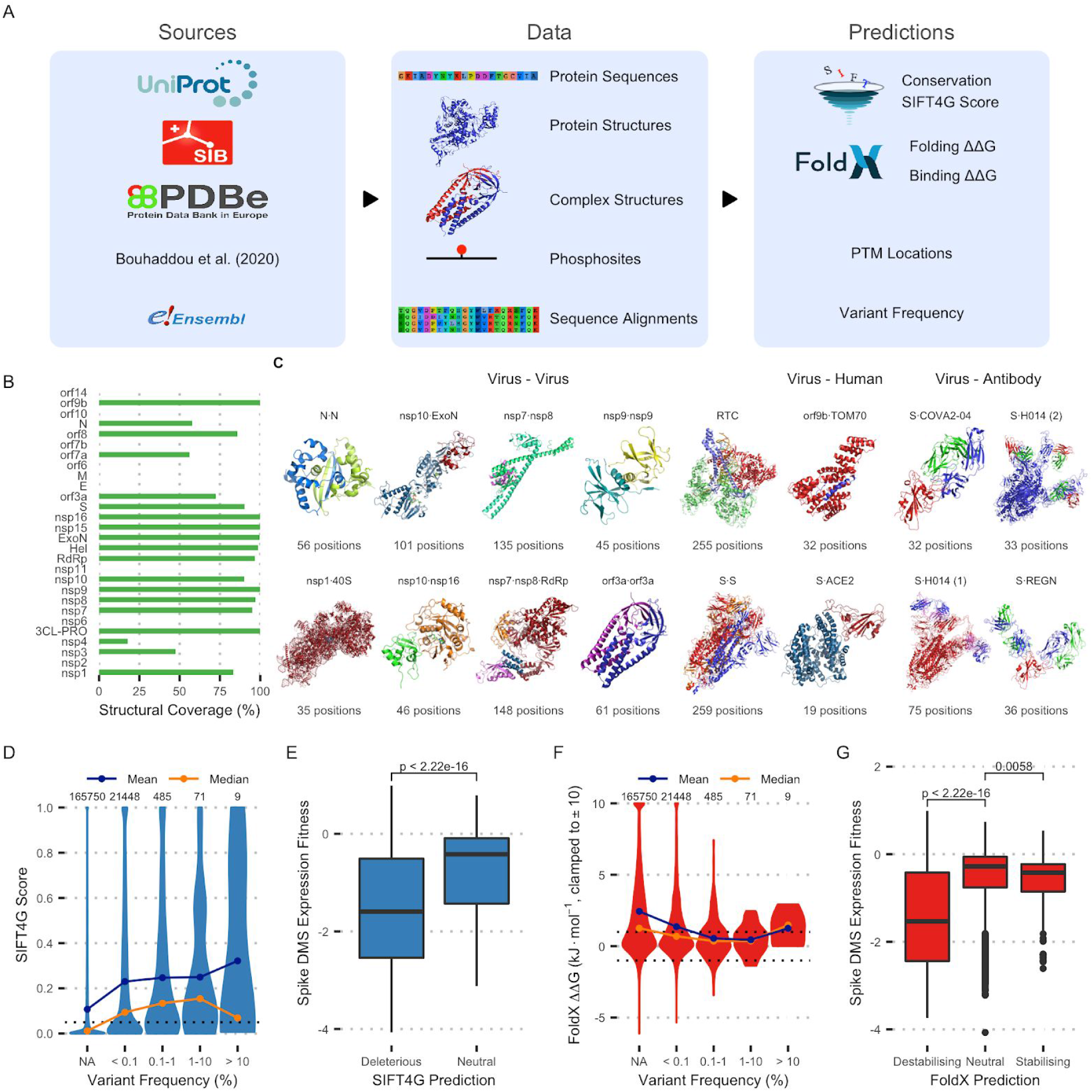
**A**: Data generation pipeline schematic. **B**: Percentage of residues covered by structural models for each protein. **C**: Protein complex structures currently included in the dataset. **D**: Distribution of SIFT4G scores for variants across frequencies. NA indicates variants that were not observed at all. The threshold for prediction being significant (0.05) is marked as is the number of variants in each category **E**: Distribution of Spike DMS variant expression fitness scores for variants predicted deleterious (< 0.05) or neutral (> 0.05) by SIFT4G. The p-value from a Wilcoxon signed-rank test is shown. **F**: Distribution of FoldX ΔΔG predictions for variants of varying frequencies. The thresholds for a variant being considered destabilising (1) and stabilising (−1) are marked. **G**: Distribution of Spike deep mutational scan variant expression fitness for variants predicted destabilising, neutral or stabilising by FoldX. P-values from Wilcoxon signed-rank tests are shown.

High frequency variants are not typically expected to impact protein function or viral fitness strongly. We used this to validate SIFT4G scores and FoldX ΔΔG predictions by comparing them to variant frequencies. In addition, we compared the predictions with the results of a deep mutational scan (DMS) experiment on Spike protein variants (Starr et al., 2020). As expected, SIFT4G scores are lower for rarer variants on average (*Figure 1D*), although there are significant deleterious predictions (< 0.05) in all frequency ranges. Variants SIFT4G predicts to be deleterious also have significantly lower fitness measurements based on the DMS experiments (*Figure 1E*). These observations support SIFT4G scores being informative for viral variants. Similar results are found for FoldX ΔΔG predictions, with higher frequency variants tending to have lower magnitude ΔΔG values (*Figure 1F*). Variants FoldX predicts are destabilising (ΔΔG > 1) have significantly lower DMS expression fitness scores (*Figure 1G*). Variants predicted stabilising (ΔΔG < −1) are not as different to neutral variants in terms of expression fitness, which is expected because stabilising proteins does not generally affect expression.

### Variant Predictions at Protein-Protein Interfaces

Many protein functions occur through interactions with other proteins, meaning accurate predictions about interactions are particularly useful. There are 1828 observed variants in the modeled interfaces, of which 443 are predicted to disturb interface binding in at least one interface (Figure 2A). We benchmarked FoldX interface binding ΔΔG in the same way as previous scores, showing higher frequency variants are predicted to be less destabilising on average (*Figure 2B*). The Spike DMS also measured the strength of binding with ACE2, one of the virus' key host targets. This is reported as a Δlog_10_K_D_, polarised so negative values mean weaker binding. Variants predicted to destabilise interface binding have significantly lower DMS binding fitnesses than variants predicted to be neutral and variants predicted to stabilise the interface have significantly higher (*Figure 2C*). Together, this suggests that the predictions reflect real effects in SARS-CoV-2 proteins and inform on variant impact.

**Figure 2.**
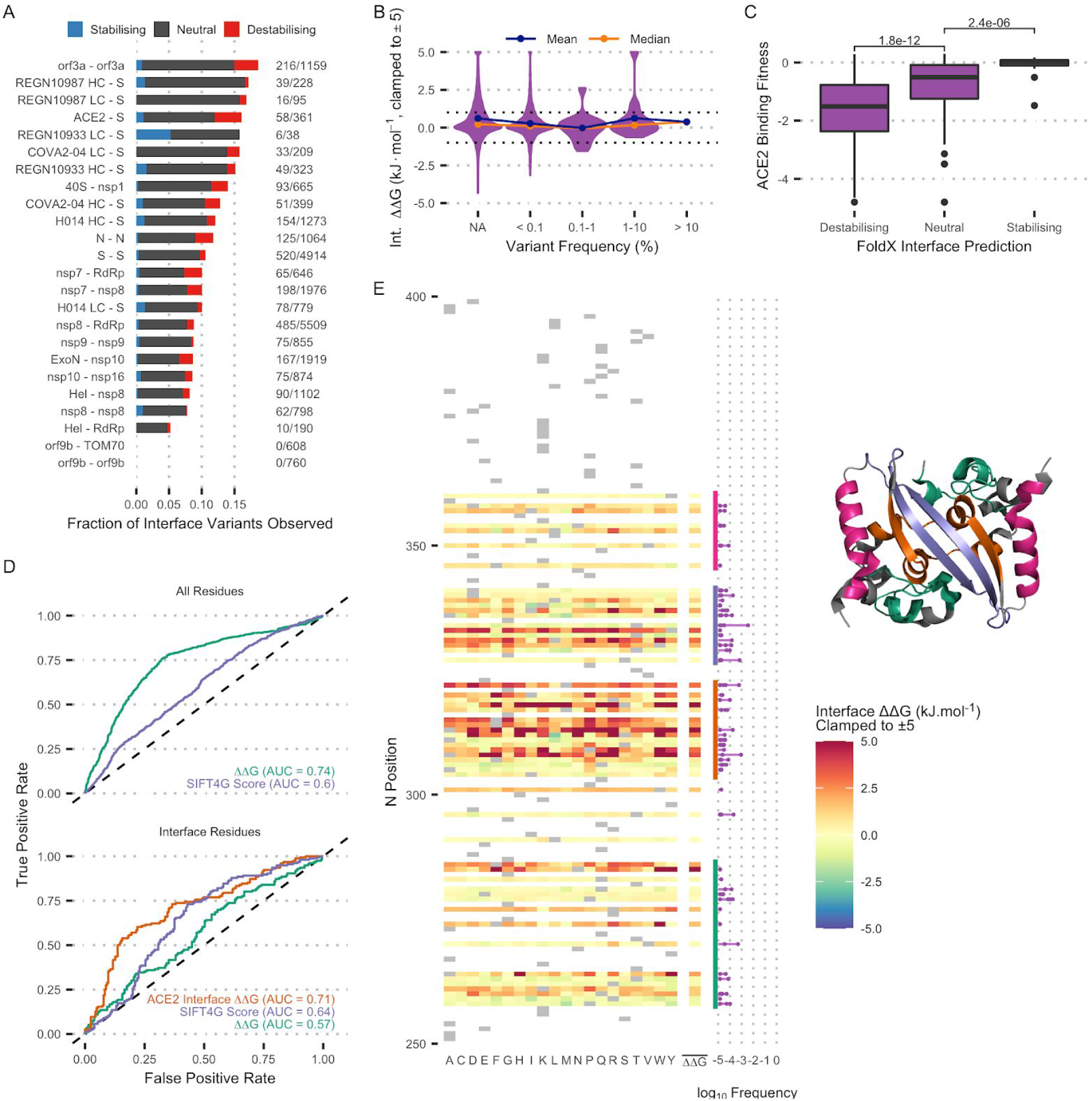
**A**: Fraction of potential interface variants observed in samples, including the fraction predicted to have significant effect. The exact number of observed and possible variants in each interface is displayed to the right. The fraction does not account for variant frequency. **B**: Distribution of FoldX’s predicted interface binding ΔΔG for variants of different frequencies. **C**: Distribution of Spike variant DMS ACE2 binding fitness for variants predicted destabilising (ΔΔG > 1), neutral (−1 < ΔΔG < 1) and stabilising (ΔΔG < −1), as well as Wilcoxon signed-rank test p-values between the groups. **D**: Sensitivity-specificity curves showing the ability of different predictors to model whether Spike variants DMS ACE2 binding fell to 10% of the WT value. The test was performed on all positions (top) and predicted interface positions (bottom). AUC is displayed for each curve. **E**: FoldX interface ΔΔG predictions for all possible variants in the N-N interface (PDBID: 7C22), mean position binding ΔΔG and the frequency of observed mutations at interface positions. The structure is also shown, coloured to indicate the marked regions in the heatmap.

We also performed a sensitivity-specificity analysis (*Figure 2D*) to assess the power of different metrics to predict whether DMS binding fitness decreased to 10% of the wild-type value, which corresponds to the long tail of deleterious results occurring in DMS scores. Analysis of interface residues shows FoldX’s interface ΔΔG for the S - ACE2 complex is the best predictor of binding changes, suggesting interface specific interactions are most important at interface residues and that this is the main metric for positions known to be in an interface. However, positions away from interfaces can also impact binding by changing overall structure. The fact that FoldX’s general ΔΔG is a good predictor when considering all residues but is weak for known interface residues suggests general structural effects are relevant for interface binding but reaffirms that interface interactions are dominant at the interface itself. SIFT4G scores perform similarly for interface and non-interface residues, suggesting conservation is a useful metric but less powerful than the appropriate structural predictions.

Predictions can also be used to analyse proteins overall properties. We visualised each interface in the dataset to identify important regions for binding and whether they are mutated. The N dimerisation interface was the most interesting of the 20 interfaces (*Figure 2E*). The two central β-strands are likely to be most important in this complex since they are where mutations cause the most destabilisation. These regions also have more observed variants than the rest of the interface, although only one at a frequency above 10^−3^ and only a small number are predicted to be destabilising themselves. This was the only case where the most impactful and most frequent interface variants co-occurred. The identification of multiple mutations within the same functional region across different strains suggests that there may be a functional advantage to mutations occuring at this interface.

### High Frequency Variants with Predicted Functional Impact

As shown above, high frequency variants tend to have lower predicted functional impacts and are typically expected to be neutral. Significant or non-neutral predicted consequences can therefore be useful to identify high frequency variants of interest. This is important when prioritising variants for follow up experiments or assessing observed variants, particularly when new strains emerge. We searched for significant predicted effects in mutations from variant strains or with elevated frequencies, highlighting a selection of examples here. In total we have identified 281 variants with frequency above 0.1% that have at least one predicted functional impact. These are provided in *Table S1* with annotations to facilitate potential follow up studies.

In December 2020 three variant strains were observed (Mahase, 2021) and have since been rapidly increasing in frequency; B.1.1.7 in the UK, B.1.351 in South Africa and P.1 in Brazil. The majority of variants in these strains are in the Spike protein (*Figure 3A*), with B.1.351 and P.1 each carrying 5 variants predicted to be structurally destabilising and B.1.1.7 carrying 3. Two variants in the Spike RBD are also observed in variant strains (all carry N501Y and E484K is in B.1.351 and P.1), both of which FoldX predicts to impact ACE2 binding. E484K, which is predicted to stabilise ACE2 binding, has also recently been observed in some B.1.1.7 samples (Public Health England, 2021). N501Y is computationally predicted to destabilise the interface but was measured to increase binding in the DMS experiment. This suggests it changes the binding conformation in a way FoldX doesn’t accurately model and emphasises that computational predictions are good at identifying variants that impact interfaces but do not always fully model the consequences. B.1.1.7 and B.1.351 also carry variants in orf8 (*Figure 3B*), which potentially interacts with the host immune system (Li et al., 2020) and vesicle trafficking (Gordon et al., 2020). The two variants in B.1.1.7 (A51I & Y73C) and the variant in B.1.351 (E92K) are all predicted to destabilise the structure. A fourth stabilising orf8 variant, S24L, occurs at a generally increased frequency (3.18%).

**Figure 3.**
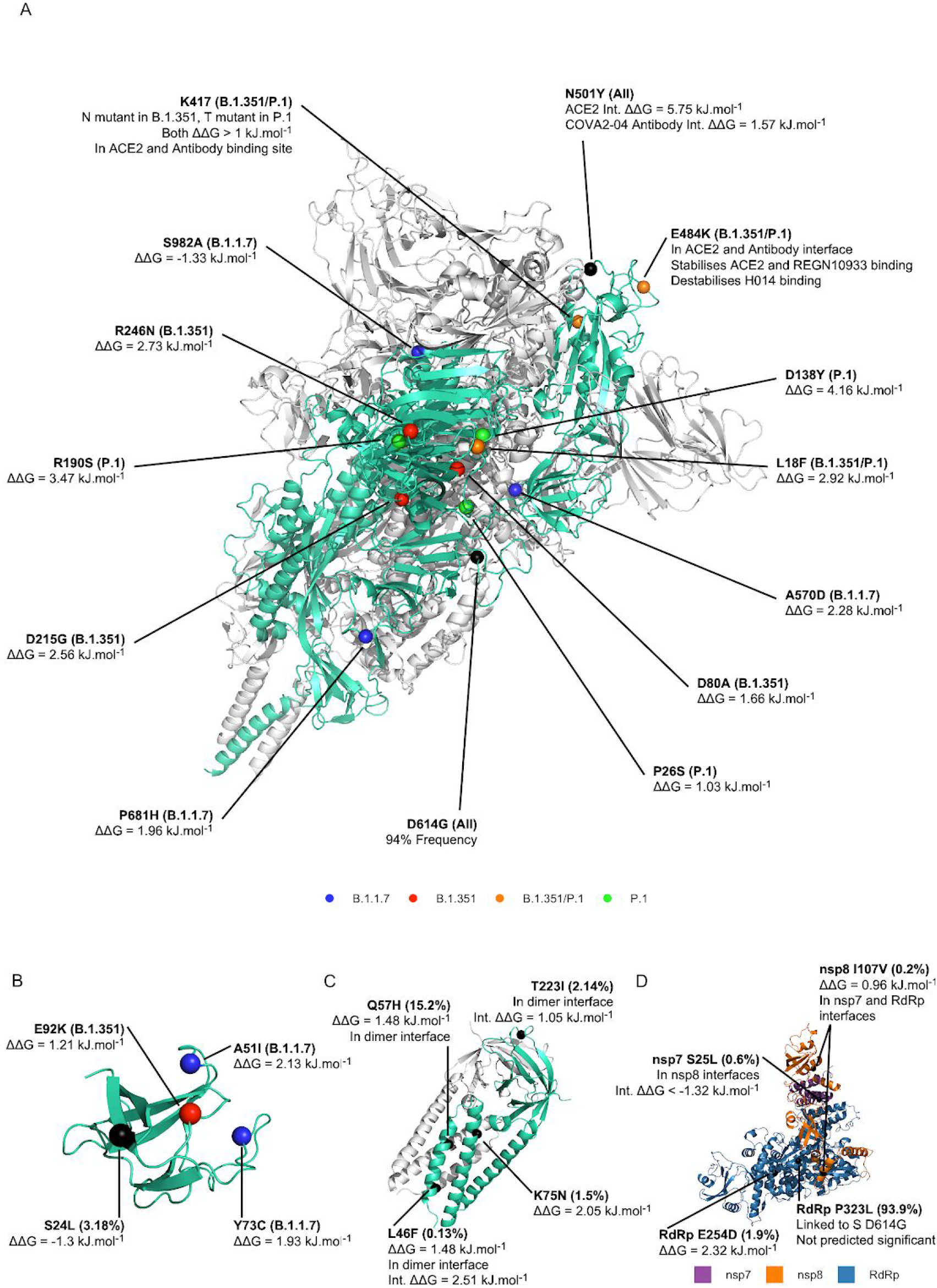
A selection of interesting observed variants and prediction details. Variants are highlighted on the structures of the Spike protein (**A**, PDBID: 6XR8), Orf8 (**B**, PDBID: 7JTL), the Orf3a homodimer complex (**C**, PDBID: 6XDC) and the nsp7 - nsp8 - RdRp portion of the replication complex (**D**, PDBID: 7BTF). Variants are coloured by variant strain where applicable.

Several other Spike variants also have elevated frequency. D614G (94%), the primary variant in the dominant B lineage, is not predicted to be significantly destabilising to either the protein or the interface, but is predicted to stop the position making contact with the other bound S subunit. A222V is known to have risen in frequency in Europe (Hodcroft et al., 2020) to 30% overall sample frequency, and is predicted to destabilise protein structure. A slightly less frequent variant, T29I (0.2%), is at a phosphosite and leads to large putative structural destabilisation.

We also highlight four variants in the orf3a ion channel dimer (*Figure 3C*), which is linked to apoptosis induction (Ren et al., 2020). Q57H (15.2%) is predicted to destabilise the protein and protrudes into the core of the transmembrane helices that form the dimerisation interface. It is not predicted to destabilise the interface but could potentially interfere with ion transport by changing from the polar neutral side chain of glutamine to the bulky basic aromatic group of histidine. L46F (0.13%) also appears to protrude into the transmembrane helix channel, with a potentially impactful change to a larger aromatic group. It is predicted to destabilise both structure and interface. T223I (2.14%) is predicted to destabilise the dimerisation interface in the cytosolic domain and K75N (1.5%) is predicted to strongly destabilise the structure.

The replication complex, responsible for replicating viral RNA, also has several variants with elevated frequencies (*Figure 3D*). The RdRp protein contains P323L (93.9%), which is linked to Spike D614G (Korber et al., 2020) but does not have any significant predicted effects. The nsp7 variant S25L (0.6%) is predicted to stabilise the interface to nsp8 in both complex models. RdRp variant E254D (1.9%) significantly destabilises the structure. Finally, nsp8 variant I107V (0.2%) is in both the nsp7 and RdRp interface and has a ΔΔG only just shy of being destabilising (0.96).

We also examined the predictions made against variants that emerged in a case study with a patient treated with convalescent antibodies (Kemp et al., 2021). The initial infection was from a strain in lineage B, carrying D614G. 26 of the 47 substitutions that rise to at least 10% frequency during the infection have at least one significant prediction associated with them (*Figure S1*). It is notable that only one sub-strain that increased in frequency did not carry at least one variant predicted to destabilise the spike protein, assuming a ΔH69/ΔV70 double deletion is destabilising.

### Predicted Impact of Antibody Evasion Variants

The ability to disturb antibody binding could have a big impact on the severity of future variant strains, especially with growing numbers being vaccinated. We tested our predictors against DMS results testing a library of Spike variants for antibody evasion potential (Greaney et al., 2021). Firstly, we tested interface predictions based on four complex structures between Spike and various antibodies (*Figure 4A*). Variants predicted to destabilise these complexes were found to be significantly more likely to experimentally evade antibody binding in 2 of 4 antibodies. This suggests the predictions reflect real effects but also illustrates the variability of antibody binding, with different antibodies responding very differently to mutations. Consequently it’s possible mutations can lead to evasion of some antibodies even if they were not measured to evade binding in the deep mutational scan.

**Figure 4.**
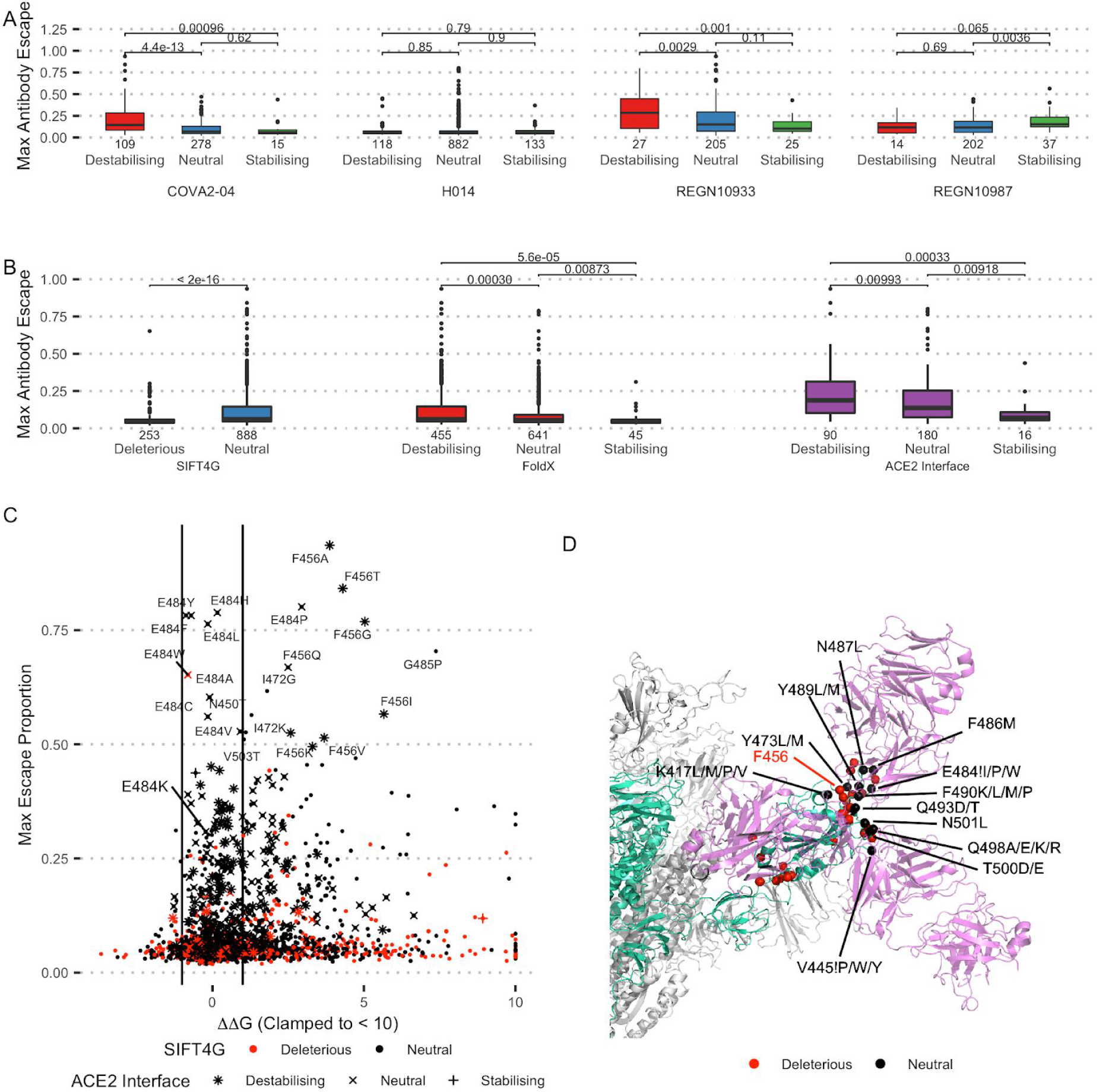
**A**: Distribution of maximum antibody evasion for variants in S - antibody complex structure interfaces, grouped by FoldX interface prediction (COVA2-04 PDBID: 7JMO, H014 PDBID: 7CAI & 7CAK, REGN10933 & REGN10987 PDBID: 6XDG). P-values from Wilcoxon signed rank tests. **B**: Distribution of maximum DMS antibody evasion scores with p-values Wilcoxon signed rank tests. **C**: Spike variant computational predictions against their maximum measured antibody evasion. FoldX ΔΔG, SIFT4G category and FoldX ACE2 interface binding indicate which variants are most likely to be selectively neutral. **D**: Variant positions with maximum evasion >10% or mean evasion >5% projected onto a Spike - antibody complex. Variants with SIFT4G Score < 0.05, |ΔΔG| > 1 or |ACE2 interface ΔΔG| > 1 are considered deleterious. All mutations to positions 445 and 484 apart from a small selection are neutral, so here the variants that are non-neutral are listed and the position marked with an exclamation mark (!). The structure is a combination of 7CAI, showing Spike (blue) in complex with H014 (pink, left), and 6XDG, which includes REGN10933 (pink, top) & REGN10987 (pink, bottom) bound to the Receptor Binding Domain. They are aligned on the RBD in order to show a variety of antibody binding configurations.

We also tested our general predictors (*Figure 4B*), since these four antibody structures do not cover all possible binding modes. Variants predicted to be destabilising by FoldX, neutral by SIFT4G or to destabilise the ACE2 - S complex binding are all significantly more likely to lead to antibody evasion. This can be understood as successful antibody evasion requiring structural changes that do not lead to loss of protein function, which makes biological sense. These results can help inform on the prevalence and emergence of Spike RBD variants with potential for antibody evasion. For instance, 41.3% of samples carry at least one Spike variant that is predicted destabilising by FoldX and neutral by SIFT4G, although of course it is likely that many of them do not interact with antibody binding. This high rate is also likely partly driven by increased sequencing of the three emergent strains, which carry such variants.

The computational predictions can also be combined with experimental antibody evasion measurements to predict which evasion variants are not predicted to be deleterious for the virus (*Figure 4C*). Variants that are predicted to be neutral based on sequence conservation, structure and ACE2 interface binding but lead to experimental antibody evasion (*Table S2)* are particularly concerning because they are less likely to be selected against. Positions leading to maximum evasion >10% or mean evasion >5% are clustered in two regions of the Spike RBD; the base of the domain connecting to the rest of the protein and the upper head, where most observed antibody interfaces occur (*Figure 4D*). Mutations in the lower cluster are all predicted to be deleterious to the protein, generally destabilising the structure and therefore more likely to impact normal protein function and be selected against. Conversely many mutations in the upper cluster are predicted to be neutral and so would be particularly concerning if observed. Of these, the two positions measured to have the greatest evasion potential are 456 and 484. No variants at 456 are predicted to be neutral but only mutation to isoleucine, proline or tryptophan are predicted to be deleterious at 484, suggesting it is one of the most important positions to monitor. E484K, which is in the emerging B.1.351 and P.1 strains and has recently been sporadically observed in B.1.1.7 (Public Health England, 2021), has a slightly lower measured evasion effect but is predicted to destabilise S - H014 antibody binding and is predicted to be neutral by SIFT4G and FoldX, reinforcing other analyses suggesting it is a particularly concerning variant (Wise, 2021). N501T, V503T, I472G and G485P are the most evasive variants at other positions, with the first two being predicted neutral and therefore of particular concern.

## Discussion

The coronavirus pandemic is a rapidly developing global health emergency, in which new mutations frequently emerge and sometimes rise to prominence. This makes the ability to rapidly assess the potential consequences of new mutations very useful. Computational predictions, alongside expert knowledge and experimentation, support this goal and help build a coherent picture of variants consequences. This is the rationale behind the Mutfunc: SARS-CoV-2 dataset, providing conservation, structure and interface based predictions alongside frequencies and phosphosite positions for all possible viral substitutions. The results are benchmarked against experimental results and frequency, showing they can aid rapid interpretation of variant impact. Our dataset complements existing SARS-CoV-2 resources, such as EVCouplings results (https://marks.hms.harvard.edu/sars-cov-2/) and the COVID3D resource (Portelli et al., 2020).

The data collated in Mutfunc: SARS-CoV-2 is a mix of computational predictions and experimental results. Frequency and phosphosite data are the results of experimental work whereas the results from FoldX and SIFT4G are computational predictions. Care must be taken when interpreting all results but particularly computational predictions. Firstly, the predictions are the result of mathematical models and have inherent uncertainty; for example FoldX ΔΔG values should not be treated like physical measurements but an indicator that a mutation is more likely to impact structure in some way, which could be beneficial or deleterious to the virus. The predictors are also not specifically predicting increases or decreases in function but predicting deleteriousness from conservation and modelling energetics. These features relate to changes in protein function and can relate to changes in important viral features such as infectivity or immune response, but are not guaranteed to. For this reason it is important to always consider specifically what a score tells you and what that means for the variant.

In addition to taking care when interpreting results it is also important to consider features that are not incorporated into the dataset. For instance, many additional interactions are experimentally observed but lack structural models (Gordon et al., 2020). This suggests there are interfaces missing, especially between virus and host proteins. The number of observed interface variants would almost certainly increase when these interactions are considered, even accounting for shared interfaces. Similarly some phosphosites were possibly not detected and other types of post-translational modifications were not considered. Other mutational consequences, for example RNA level effects, are not considered at all, despite potential functional consequences.

Despite these cautions, the dataset of predicted variant effects allowed us to search observed variants and identify those that appear most likely to impact protein function, based on frequency, computational predictions and our knowledge of the proteins. This analysis identified various variants that have already been discussed in the literature, as well as other potentially interesting uninvestigated variants. A similar analysis combining predictions with experimental antibody evasion measurements allowed us to identify variants most likely to cause antibody evasion and not negatively impact viral fitness. Such results enhance monitoring by highlighting potentially important variants for deeper analysis and identifying variants to monitor.

The dataset is available to download and search at sars.mutfunc.com. We hope that this resource will aid researchers assessing the impacts of viral variants, complementing and informing experiments and expert analyses.

## Methods

The code managing the pipeline and analyses are available at https://github.com/allydunham/mutfunc_sars_cov_2. The web service source code is available at https://github.com/allydunham/mutfunc_sars_cov_2_frontend. The pipeline is managed through Snakemake (Köster and Rahmann, 2012).

### Conservation

SARS-CoV-2 protein sequences were downloaded from Uniprot (The UniProt Consortium, 2019) and the orf1ab polyprotein split into sub-sequences based on the Uniprot annotation. A custom reference database was generated based on the NCBI virus coronavirus genomes dataset (NCBI Resource Coordinators, 2018), which includes sequences from a large range of coronaviruses. SARS-CoV-2, SARS and MERS sequences were filtered to only contain sequences from the Wuhan-Hu-1 strain, the Urbani strain and the HCoV-EMC/2012 strain respectively. Without this the dataset contains very large numbers of almost identical sequences from patient samples, which are not informative since SIFT4G looks to compare across species. The remaining sequences were clustered using MMseqs2 (Steinegger and Söding, 2017) with an overlap threshold of 0.8 and a sequence identity threshold of 0.95, which grouped other duplicate sequences into representative clusters. SIFT4G Scores were generated for all possible variants to the SARS-CoV-2 sequences based on this database. A modified copy of SIFT4G was used, which reports scores to 5 decimal places instead of the usual 2.

### Structural Destabilisation

Structures were sourced from the SWISS-Model (Bienert et al., 2017; Waterhouse et al., 2018) SARS-CoV-2 repository (https://swissmodel.expasy.org/repository/species/2697049), which contains experimental structures and homology models. Models were required to have greater than 30% sequence identity and a QMean score (Benkert et al., 2011) greater than −4, as recommended by SWISS-Model. Suitable models were available for 19 of the 28 viral proteins. Models were ordered by priority; firstly experimental models over homology models and then by QMean Score. Models were examined in turn and any position not covered by a higher priority model was added to the FoldX analysis pipeline. FoldX's RepairPDB command was used to pre-process selected SWISS-Model PDB files. All mutations at each position assigned to each model were modeled using the BuildModel command, using the average ΔΔG prediction from three runs.

### Surface Accessibility

Naccess (Hubbard and Thornton, 1993) was run on each structure using the default settings. Structures were filtered to only include the chain corresponding to the appropriate SARS-CoV-2 protein. This means some surface accessible positions are usually found in interfaces rather than facing the solvent. Since structures are not always complete surface accessibility is an approximation and will not be accurate in all cases.

### Protein Interfaces

Complex models were identified from SWISS-Model, literature search and PDBe (Gordon et al., 2020; Henderson et al., 2020; PDBe-KB Consortium et al., 2020; Schubert et al., 2020; Zhou et al., 2020). PDB files were pre-processed using the FoldX RepairPDB command. An initial AnalyseComplex command was used to identify positions involved in each interface and estimate the energetics of the wild-type interface. Structural models of all single amino acid substitutions to the interface were generated using the BuildModel command and the mutant interfaces re-analysed with AnalyseComplex. The wild-type energetic predictions were subtracted from the mutants’ to estimate energetic changes and if any amino acids had been lost or gained from the interface.

### Variant Frequency

Frequencies are based on an alignment containing observed mutations from public SARS-CoV-2 sequences from COG-UK, GENBANK and The China National Center for Bioinformation, derived from that used for sarscov2phylo. It was filtered to exclude problematic sites using VCFTools, based on the annotation at https://github.com/W-L/ProblematicSites_SARS-CoV2/blob/master/problematic_sites_sarsCov2.vcf. Variants marked as seq_end, ambiguous, highly_ambiguous, interspecific_contamination, nanopore_adapter, narrow_src or single_src were excluded because of high potential error rates. VCFTools was used to calculate frequencies, including frequencies based on regional and recent subsets of the samples. The SARS-CoV-2 genome was sourced from Ensembl (Yates et al., 2020) and Tabix indexed. Variants were annotated to genes using the Ensembl VEP tool using a custom annotation based on Ensembl’s annotation but with polyproteins split into sub-regions so that VEP assigned variants correctly. Only coding variants were considered.

## Supplementary

**Figure S1.**
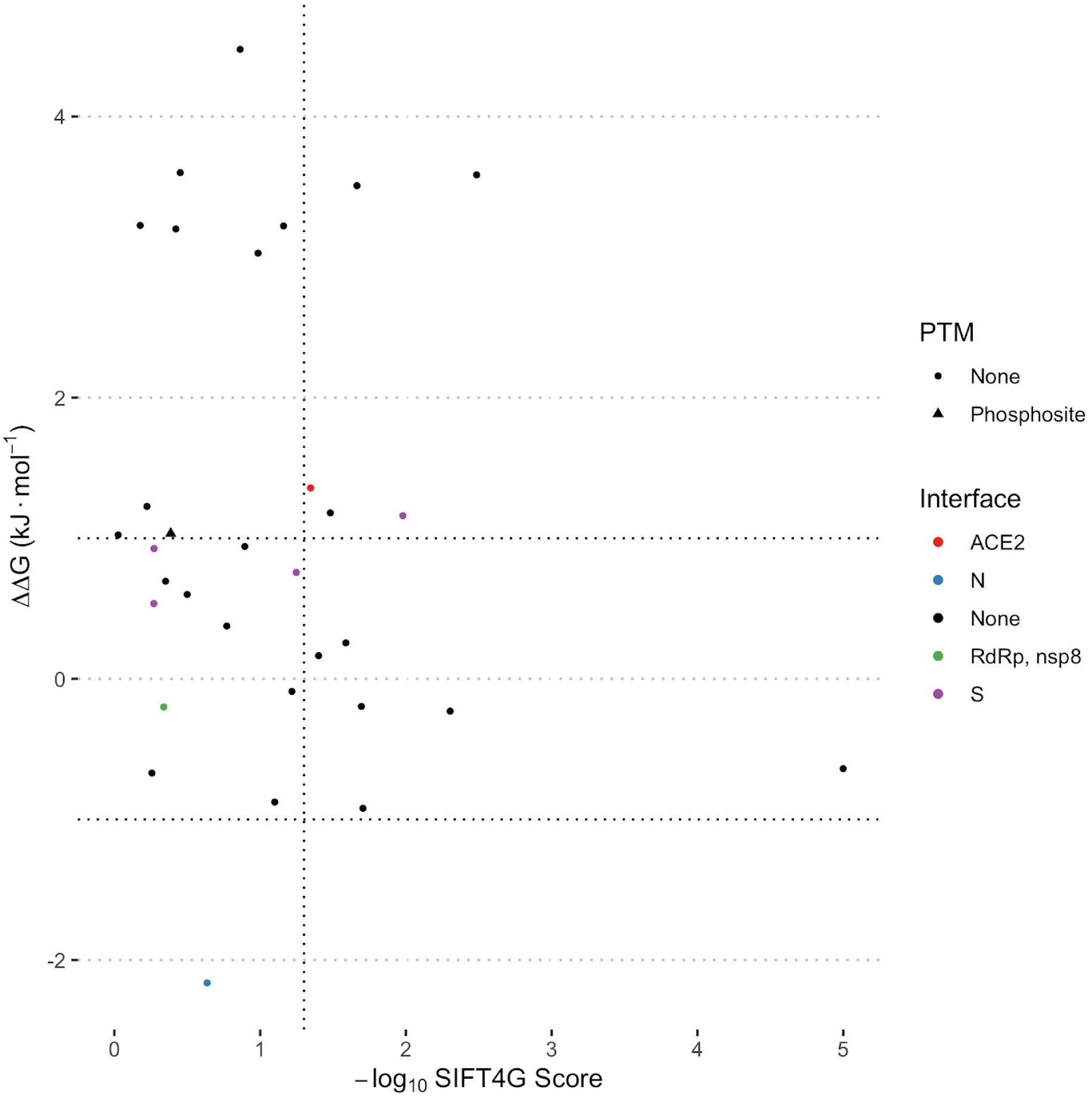
Computational predictions for variants observed to increase to at least 10% frequency at some point during the case study observed by Kemp et al. (2021).

## Notes

http://sars.mutfunc.com

### Competing Interest Statement

The authors have declared no competing interest.

